# Evaluation of LiDAR-based Canopy Trait Estimation in Midwestern Row Crops

**DOI:** 10.1101/2025.09.28.678542

**Authors:** Matthew H Siebers, Peng Fu, Stephen P Long, Justin M McGrath, Carl J Bernacchi

## Abstract

Stand count, the number of plants per unit ground area, and leaf area index (LAI), the ratio of leaf area to ground area, are critical traits for crop research but are traditionally measured using labor-intensive methods. While new sensing technologies are being developed, quantifying improvement in measurement efficiency and data quality, relative to traditional techniques, is lacking. In this study, we use LiDAR to generate 3D scans of corn and soybean plots and evaluate two computational methods: a gap fraction approach to estimate LAI and a persistent homology algorithm to estimate stand count by detecting structural peaks in the canopy. Validation experiments and statistical comparisons of bias and variance demonstrate that LiDAR-derived LAI estimates in corn are comparable in quality to those from established instruments. However, in soybean, the LiDAR method performs poorly, likely due to dense canopies limiting light penetration and structural differentiation. Stand count estimations in corn closely match manual counts, with the added benefit of full-plot coverage and significantly faster data collection. In soybean, stand count estimates are unreliable under dense canopy conditions. These results offer practical guidance for the use of LiDAR in field phenotyping and highlight both its current capabilities and limitations. While a trade-off between speed and precision remains, particularly in high-density canopies, LiDAR’s scalability and multi-trait potential make it a promising tool for high-throughput breeding programs. Continued improvements in LiDAR hardware and algorithm design may further enhance measurement accuracy and extend applicability across crops and growth stages.

## Introduction

Phenomics tools have the potential to accelerate crop breeding by increasing the number of measurable traits across more crop varieties than traditional methods allow (Zhao et al., 2019). Realizing this potential requires developing and validating new capabilities to produce high-quality data tailored to plant breeders’ needs (Deery & Jones, 2021). This process typically involves selecting a sensor, designing trait-estimation algorithms, and validating those estimates against established methods. Stand count, the number of plants per unit row length, is valuable for interpreting plot-level biomass variation, helping disentangle the confounding influence of plant number vs. plant performance (Shi et al., 2015). Stand count can also aid in interpreting crop stress responses; for example, salt tolerance may be evaluated by counting plants that germinate under saline conditions (Ahmed et al., 2024). Similarly, Leaf Area Index (LAI), the ratio of leaf area to ground area (m^2^/m^2^), is crucial for breeders and ecosystem modelers. Key physiological processes such as transpiration, light interception, and photosynthesis are directly influenced by LAI (Huang et al., 2015; Lochocki et al., 2022; Morakinyo & Lam, 2016).

Traditional methods for determining stand count and LAI are time-consuming, limiting measurements across space (e.g., total number of plots) and time (frequency of measurements). Traditionally, stand count is obtained manually by counting plants over a known row length. The gold-standard method for LAI involves harvesting all plants in a row segment, removing and measuring the leaves, and calculating LAI from total leaf area. Less destructive approaches use hemispherical photography or canopy analyzers which estimate LAI based on visible sky within the canopy using established light-penetration models. Though implementation varies, most require several in-canopy readings and subsequent processing. These methods are faster than harvesting but still labor-intensive, taking several minutes per measurement (Bréda, 2003). Since stand count and LAI are architectural traits, accurate three-dimensional (3D) reconstruction of the plant canopy could enable their computational estimation. Light Detection and Ranging (LiDAR) offers a promising high-throughput solution to 3D canopy imaging and is widely used in plant phenomics to study growth and structure (Bailey & Mahaffee, 2017; D. Deery et al., 2014; Jimenez-Berni et al., 2018; Madec et al., 2017; Siebers et al., 2018, 2024). LiDAR emits light pulses that reflect off surfaces and return to the sensor; by measuring time of flight relative to sensor position, detailed 3D point clouds are generated (Rosell et al., 2009). While these point clouds capture plant structure, they do not inherently include trait data (Rosell Polo et al., 2009/2), necessitating specialized algorithms to extract meaningful information.

Persistent homology is well-suited for identifying geometric features in 3D data (Edelsbrunner & Harer, 2008). It not only detects the most prominent or densest peaks but also evaluates their persistence relative to surrounding peaks, enabling it to filter out noise. Thus, it offers a novel approach for estimating stand count. As persistent homology algorithms have not previously been used for counting plants in LiDAR data, rigorous analysis is necessary to assess and validate its performance. Additionally, several methods have been compared for determining LAI, with previous studies having demonstrated agreement among several traditional measurement techniques, with each exhibiting design-specific tradeoffs (Wilhelm et al., 2000). LAI estimation using LiDAR coupled with a gap fraction algorithm has been proposed previously (McGrath et al., 2023), however it has not yet been evaluated alongside traditional measurement techniques.

This study uses LiDAR technology to estimate stand count and LAI in two major crops, corn (*Zea mays*) and soybean (*Glycine max*), aiming to statistically compare these methods with conventional ones. We apply a persistent homology algorithm, providing a novel method for estimating stand count from LiDAR data. Stand count is derived from LiDAR point clouds by detecting individual plants as clusters of points. The cloud is divided into vertical columns, and the number of points per column is used to create a two-dimensional density map. Peaks in this map, which indicate areas of higher point concentration, correspond to plant stems. This approach simulates a bird’s-eye view of the canopy overlaid with a grid to identify dense regions. Our analysis also provides updated bias and variance comparisons of LAI, which overcome the limitations with traditional comparative statistics (Goodwin & Leech, 2006; Taffé, 2021). We predict that LiDAR-based methods will perform best in canopies with clearly separated plants and well-defined gaps. This research aims to advance the utility of LiDAR-based measurements of LAI and stand count in plant phenotyping and to advance adoption of these techniques in breeding and agronomic applications.

## Methods

### 2.1 Field Sites, Experimental Design, and Data Collection

The experiment consisted of 12 soybean plots (*Glycine max* var. LD11-2070) and 15 corn plots (*Zea mays* var. DKC58 34RI) that were planted on June 7, 2023. Each plot consisted of four rows that were 3.05 m (10 ft) long. Row spacing for soybean and corn plots was76 cm (30 inch). On August 16, 2023, nine of the 15 maize plots were thinned manually to create three plots each with 25%, 50%, and 75% of the plants removed, and six unthinned control plots with maximum planting density. A similar thinning treatment was applied for soybean on September 8, 2023, with three control plots at maximal planting density.

Stand count and LAI were measured multiple times per plot on August 21 for corn and September 21 for soybean to assess method variance. Five trained individuals independently assessed stand count using three methods: meter sticks, fast-paced LiDAR scans, and slow-paced LiDAR scans. Each method was applied five times per plot. “Fast pace” refers to a typical walking speed while pulling the cart (∼0.5 to 1 m/s), and a “slow pace” was less than 0.5 m/s. The meter stick method relied on a one-meter ruler placed parallel to the two center rows at a random distance from the front edge of the plot. All plants along the 1-meter transect were counted, with one end of the transect considered inclusive and the other end exclusive to reduce edge-effect bias.

LAI was measured using four instruments: the LAI-2200 (LI-COR Environmental, Lincoln, NE, USA), AccuPAR (Meter Group, Pullman, WA, USA), SunScan (Delta-T Devices Ltd, Cambridge, UK), and both fast and slow LiDAR scans. LiDAR-based LAI was estimated using gap fraction analysis (McGrath et al., 2024). Five people were trained to collect LAI measurements in the thinned corn and soybean plots. Time stamps from the instrument data files were used to estimate how long each person took to complete all measurements for each crop. The same LiDAR scans were used for stand count and LAI estimation.

LiDAR point clouds were collected using a 2D LiDAR sensor (Hokuyo UST-10LX) and a rotary encoder (Yumo E6B2-CWZ3E) mounted on a modified garden cart (Gorilla Carts, Eden Prairie, MN, USA; Fig. 1). Data from the LiDAR and encoder were wirelessly transmitted to a laptop and stored for processing (Siebers et al., 2024). Point clouds were generated by synchronizing time stamps from both devices to calculate x, y, and z coordinates for each point.

**Figure 1.**
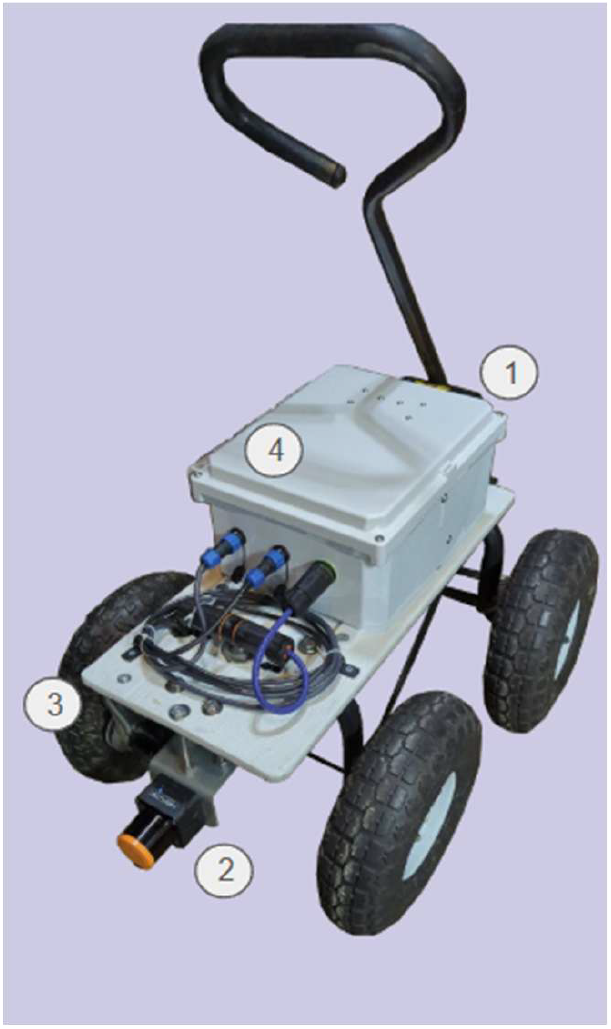
A lidar-based 3D scanning system. The system was powered by a 20 V lithium-ion battery (1). A 2D scanning LiDAR (Hokuyo LX-10, Hokuyo Automatic Co., Ltd., Osaka, Japan) was mounted to the cart 7 inches above the ground (2). The distance traveled was measured using a rotary encoder (3, E6B2-CWZ3E, Yumo automation, Yueqing China) connected to the wheel with a timing belt. A wireless access point (TP-Link EAP 110, TP-Link Technologies Co., Ltd., Shenzhen, China) was housed in an enclosure. The access point transmitted data which were stored on a laptop, wirelessly connected to the cart.

### 2.2 Algorithm Application

LAI was calculated from the LiDAR data following the approach of McGrath et al. (2023). Briefly, after filtering, the fraction of unobstructed angles in defined zenith intervals was used to estimate canopy openness. These gap fractions were then converted into LAI estimates, with adjustments made for cases where no gaps were observed in a given interval.

A simplified version of the persistent homology algorithm was applied to detect peaks in point density, following the method described by Huber (2021). The two-dimensional point density for each LiDAR scan was determined by dividing the x–z plane (parallel to the ground) into an irregular grid. Along the x-axis, grid cells were 2 cm wide; along the z-axis, each individual scan formed a cell. Although LiDAR scans at regular intervals, variations in cart speed result in uneven spacing along the z-axis. This grid defines vertical columns in the x–z plane, and density was calculated by counting the number of points within each column.

The persistent homology algorithm assigns each peak a “birth” and “death” height. The birth height represents the initial appearance of a peak, while the death height marks when it merges with a taller neighboring peak. This process can be visualized as submerged mountains in a sea with receding water levels. As the water level drops, individual peaks emerge, representing the birth height. As the water continues to recede, some peaks merge with taller peaks at their death height. The persistence of a peak, the difference between its birth and death heights, measures how long it remained distinct and thus how topologically significant it is in the data set.

Areas around plant stems include points from both the stem and the surrounding leaves. These produce tall density peaks with high persistence, as they persist until merging with neighboring plant peaks. In contrast, small density peaks from leaves also appear tall but exhibit low persistence due to early merging. By selecting only peaks with high persistence, those associated with individual plants can be isolated. The maximum height in each scan was normalized to 1, and peaks with persistence greater than 0.35 were counted as individual plants. Because the x–z coordinates of each peak are known, their locations can be visualized in the 3D point cloud.

### 2.3 Statistical analysis of bias and variance

The mean of all measurements was calculated for each crop, plot, and measurement method. Bias between methods was defined as the difference in mean values for each pair of methods. To evaluate bias, these differences were plotted against the mean of both methods for each plot. Plots were visually inspected to determine whether bias appeared to correlate with the mean. If no such correlation was observed, biases were pooled, and an overall mean bias was tested for significant deviation from zero using a two-tailed, two-sample t-test with a significance level of α = 0.05.

For each plot and method, the standard deviation of the repeated measurements was calculated and plotted against the corresponding mean. These plots were visually assessed for correlation between standard deviation and mean. In all cases, a correlation was observed, indicating that variances could not be pooled across plots. Consequently, for each plot, the ratio of variances between methods was tested for significant deviation from one using a two-tailed F-test at α = 0.05.

## Results

### 3.1 Persistent Homology Estimations of Plant Number Correlated Well with Manual Counts

LiDAR and encoder data were combined to produce 3D point clouds of plants (Fig. 2). In corn, both fast- and slow-paced LiDAR-derived stand count estimates correlated strongly with manual counts (Fig. 3A: fast, *r* = 0.92; Fig. 3D: slow, *r* = 0.93). The bias between LiDAR and manual estimates increased with stand density, with undercounting most pronounced in plots with more than four plants per linear meter (Fig. 3B, E). None of the fast LiDAR estimates were significantly more variable than manual counts. In contrast, four hand-counted plots showed significantly greater variance than the corresponding slow LiDAR estimates (Fig. 3F).

**Figure 2.**
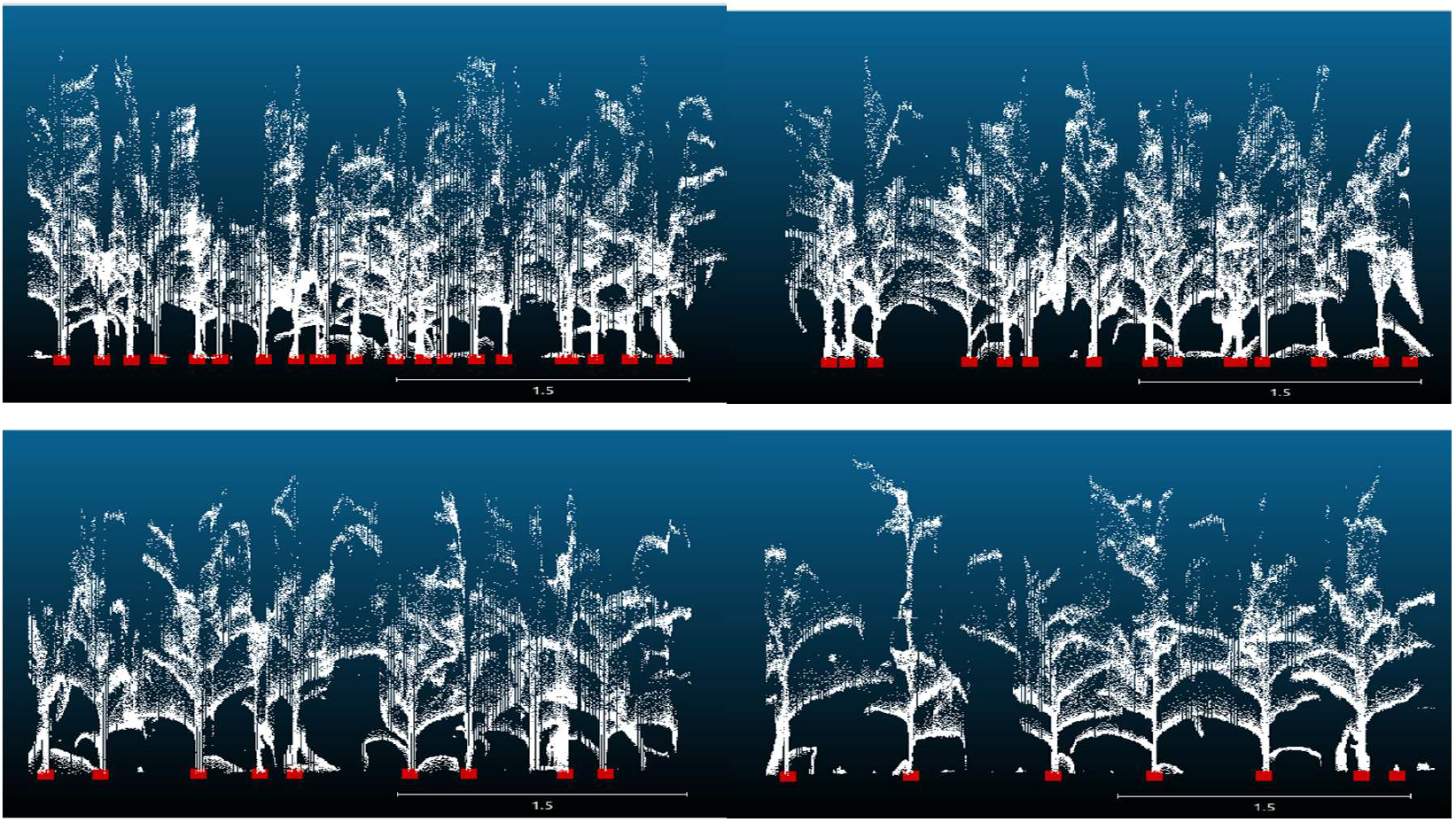
Example LiDAR scan showing one row from corn plots thinned at 0% (a), 25% (b), 50% (c), and 75% (d). Red points at the base of each image indicate peak locations identified by the persistent homology algorithm. Scale bar shown in meters.

**Figure 3.**
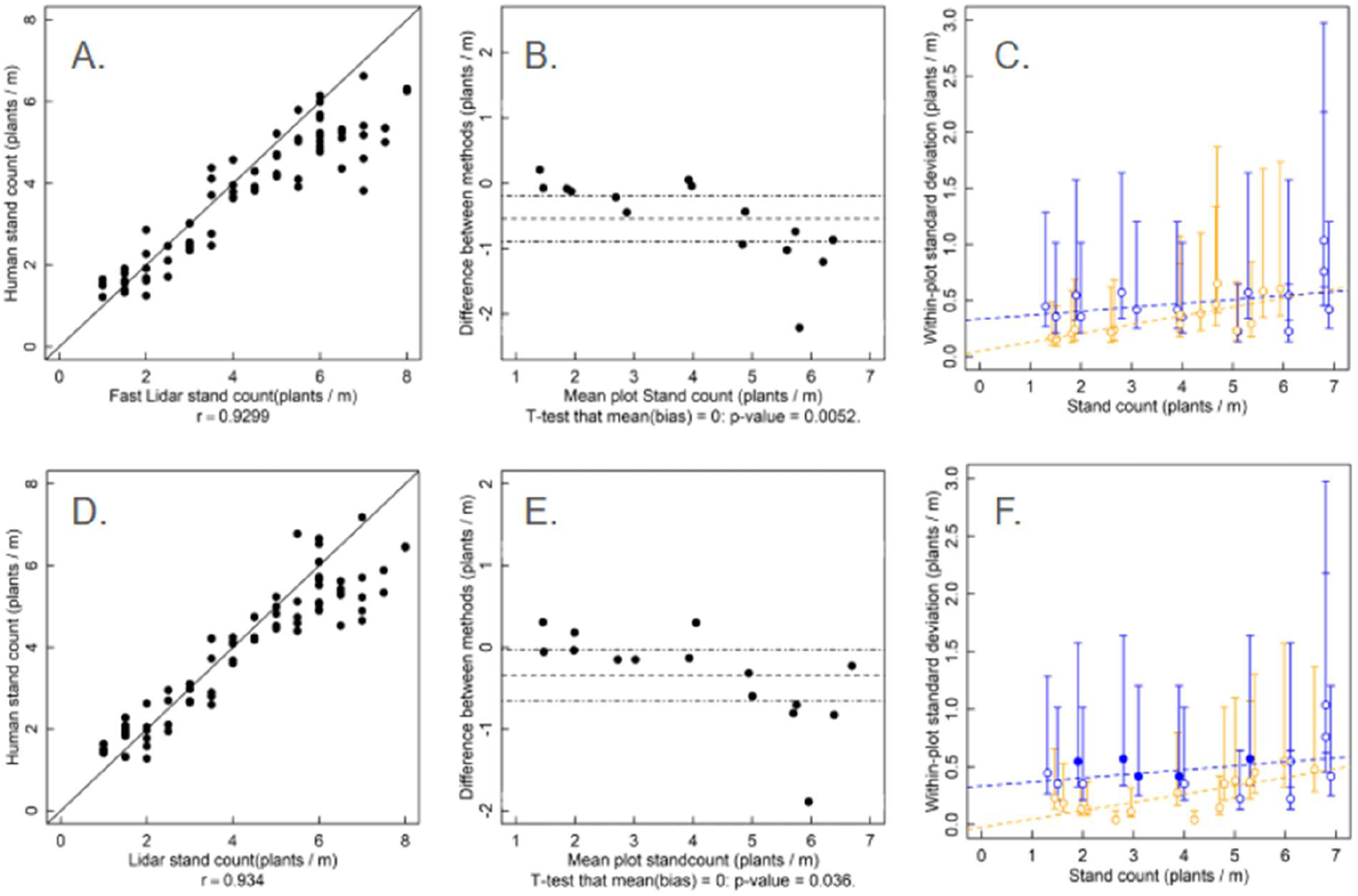
Correlation (A, D), bias (B, E), and standard deviation (C, F) between hand-counted and LiDAR-estimated stand counts (plants/m) in corn. Data were collected from 15 plots thinned at 0%, 25%, 50%, and 75%. Bias plots show the mean of five repeated measurements per method (dots), mean bias (dashed line), and 95% confidence limits of the bias (dashed-dotted line). Standard deviation plots compare manual counts (blue symbols) with LiDAR estimates (orange symbols) using two-tailed F-tests. Filled symbols indicate significantly greater variance in that method compared to the other method within the same plot (*p* < 0.05). Dashed lines in variance plots are least-squares regressions shown for visualization only.

In soybean, persistent-homology-based estimates correlated poorly with manual counts for both fast (*r* = 0.59; Fig. 4A) and slow (*r* = 0.58; Fig. 4D) LiDAR scans. Estimates were biased, with bias increasing at higher plant densities (Fig. 4B, E). Fast and slow LiDAR scans were not significantly less precise than manual counts (Fig. 4C), and slow scans were equally precise (Fig. 4F). Overall, the algorithm failed to accurately detect individual soybean plants, returning similar stand counts regardless of actual plant number.

**Figure 4.**
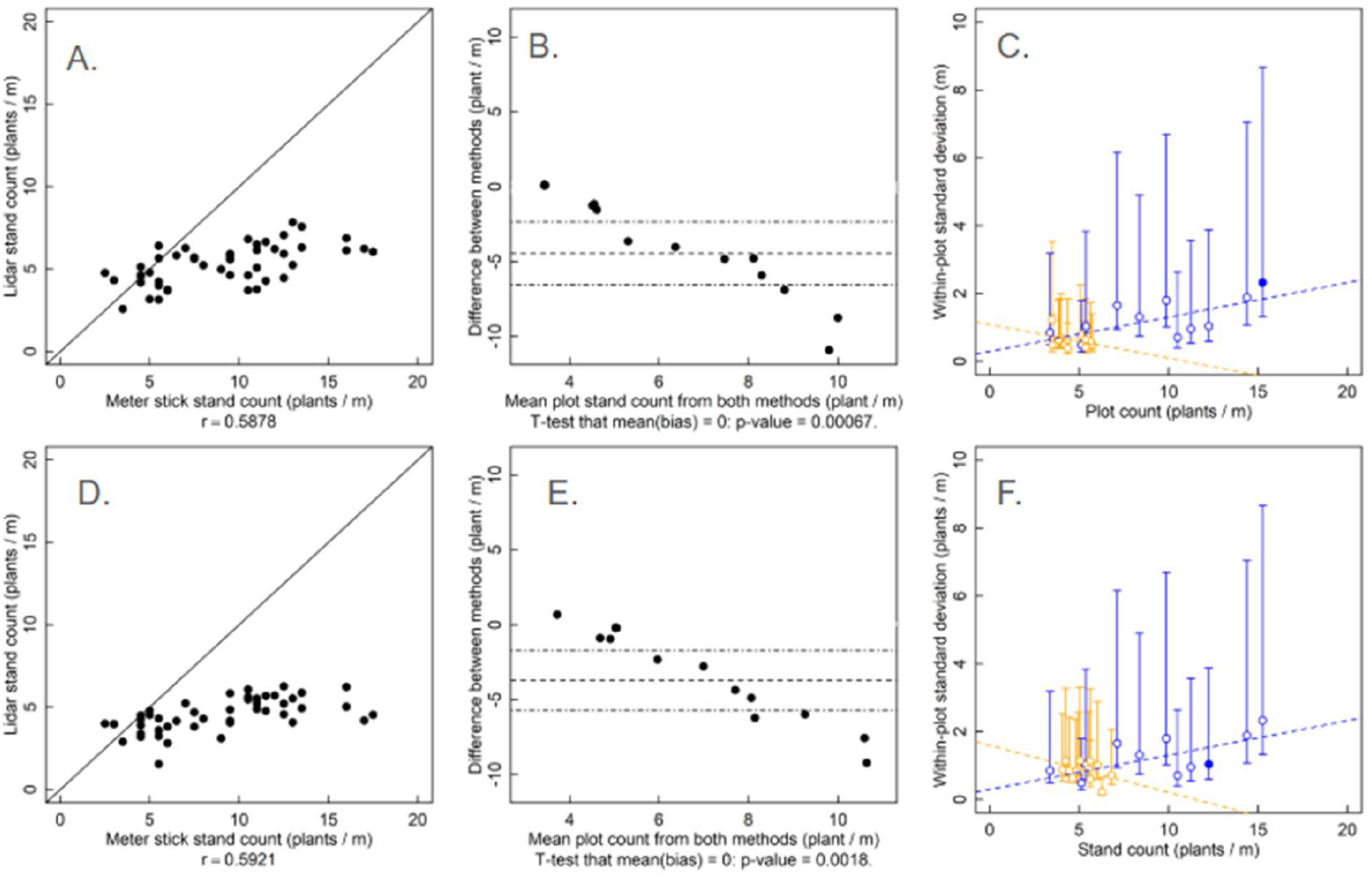
Correlation (A, D), bias (B, E), and standard deviation (C, F) between hand-counted and LiDAR-estimated stand counts (plants/m) in soybean. Data were collected from 12 plots thinned at 0%, 25%, 50%, and 75%. Bias plots show the mean of five repeated measurements per method (dots), mean bias (dashed line), and 95% confidence interval of the bias (dashed-dotted line). Standard deviation plots compare manual counts (blue symbols) with LiDAR estimates (yellow symbols) using two-tailed F-tests. Filled symbols indicate significantly greater variance in that method compared to the other within the same plot (*p* < 0.05). Dashed lines in panels C and F are least-squares regressions shown for visualization only.

### 3.2 Leaf Area Index

In corn, all measurement methods correlated well with the LAI-2200, with *r* values exceeding 0.8 (Table 1). The AccuPAR was the only instrument biased relative to the LAI-2200, underestimating LAI by an average of 0.25 units (Supplemental Fig. 1). The SunScan and both LiDAR scan speeds were more precise than the LAI-2200 at low LAI values. LiDAR scanning was the fastest method overall: fast scans required about one-quarter of the time, and slow scans less than half the time, of that needed to complete LAI-2200 measurements (Table 1).

**Table 1.**
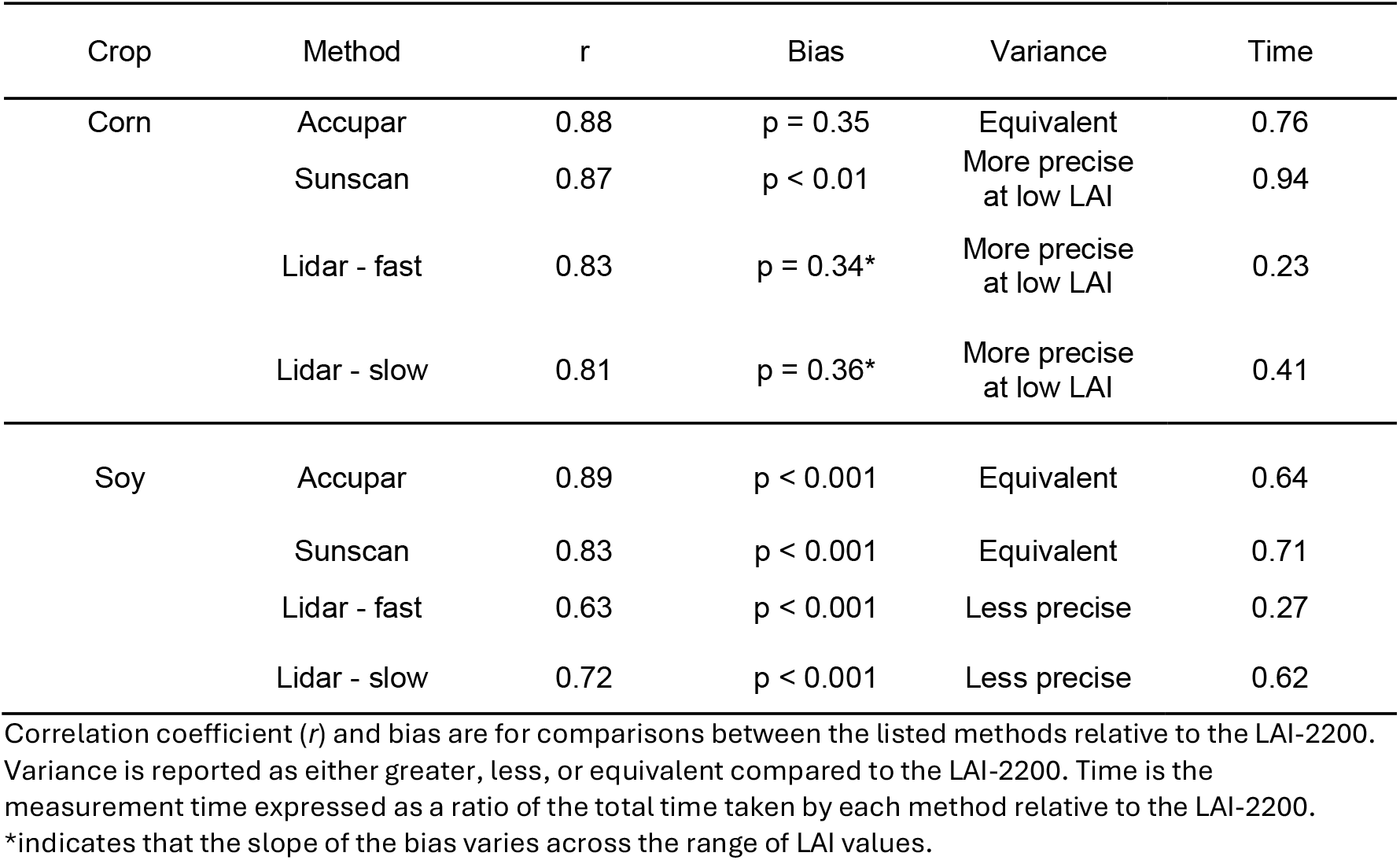
Summary of statistical comparisons between LAI measurement methods and the LAI-2200. Each method is evaluated for bias and variance relative to the LAI-2200.

| Crop | Method | r | Bias | Variance | Time |
| --- | --- | --- | --- | --- | --- |
| Corn | Accupar | 0.88 | p = 0.35 | Equivalent | 0.76 |
|  | Sunscan | 0.87 | p < 0.01 | More precise at low LAI | 0.94 |
|  | Lidar - fast | 0.83 | p = 0.34* | More precise at low LAI | 0.23 |
|  | Lidar - slow | 0.81 | p = 0.36* | More precise at low LAI | 0.41 |
| Soy | Accupar | 0.89 | p < 0.001 | Equivalent | 0.64 |
|  | Sunscan | 0.83 | p < 0.001 | Equivalent | 0.71 |
|  | Lidar - fast | 0.63 | p < 0.001 | Less precise | 0.27 |
|  | Lidar - slow | 0.72 | p < 0.001 | Less precise | 0.62 |

Correlation coefficient (*r*) and bias are for comparisons between the listed methods relative to the LAI-2200. Variance is reported as either greater, less, or equivalent compared to the LAI-2200. Time is the measurement time expressed as a ratio of the total time taken by each method relative to the LAI-2200.
\*indicates that the slope of the bias varies across the range of LAI values.

In soybean, SunScan and AccuPAR estimates also correlated well with those from the LAI-2200 (Table 1). However, all instruments showed bias compared to the LAI-2200. SunScan was less variable than the LAI-2200 in two of the 12 plots, and AccuPAR less variable in one plot. As with corn, LiDAR scanning was the fastest method, with fast scans requiring roughly a quarter of the time, and slow scans less than half the time, of that needed for the LAI-2200 (Table 1).

In corn, the methods differed in both their central tendency and spread across the thinning treatments (Fig. 5). In corn, AccuPAR was more precise than the LAI-2200 in plots with 75% thinning (*p* = 0.03) but had equivalent precision at other thinning levels. Fast and slow LiDAR scans were less precise than the LAI-2200 in control plots (*p* = 0.004 and *p* = 0.002, respectively) and more precise in plots with 50% thinning (*p* = 0.01 and *p* = 0.02; Fig. 5).

**Figure 5.**
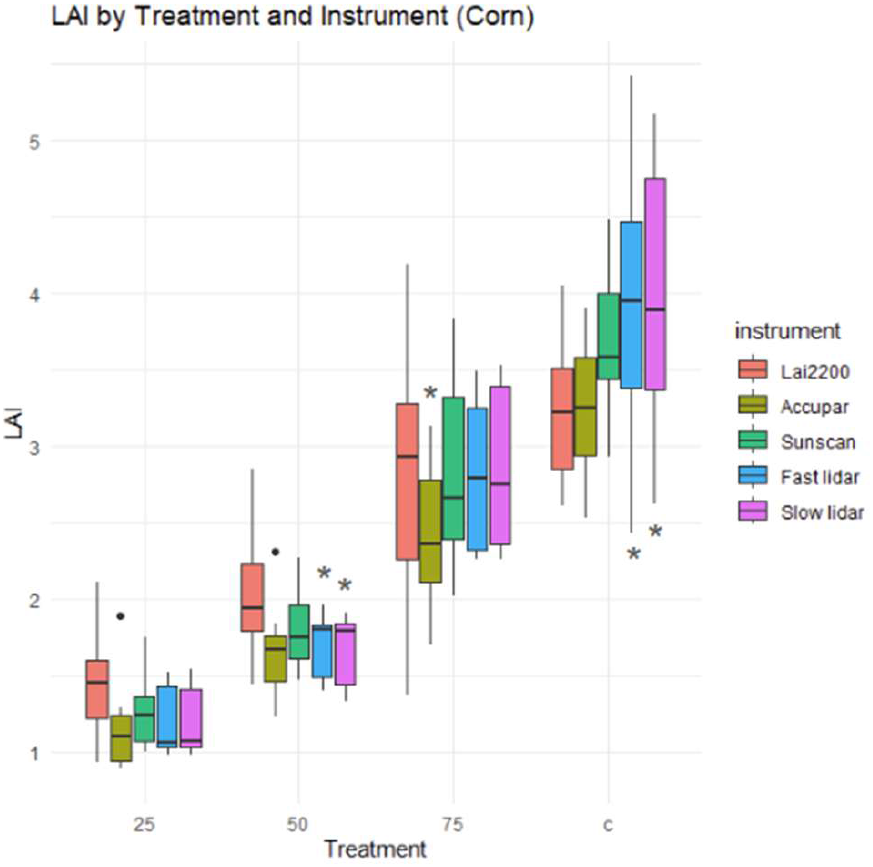
Distributions of LAI measurements in corn for each method—AccuPAR, LAI-2200, SunScan, and fast and slow LiDAR—across thinning treatment levels: control, 75%, 50%, and 25% of initial plant density. Each boxplot includes all repeated measurements for a given instrument within a treatment. At each treatment level, variance in LAI for each method was compared to that of the LAI-2200 using a two-tailed F-test. Significant differences in variance are indicated with an asterisk (*p* < 0.05).

In soybean, all LAI meters were biased relative to the LAI-2200 (Table 1; Fig. 6; Supplemental Fig. 2). SunScan was more precise than the LAI-2200 in control plots and plots with 75% thinning. Both LiDAR scanning speeds were less precise than the LAI-2200 in control plots.

**Figure 6.**
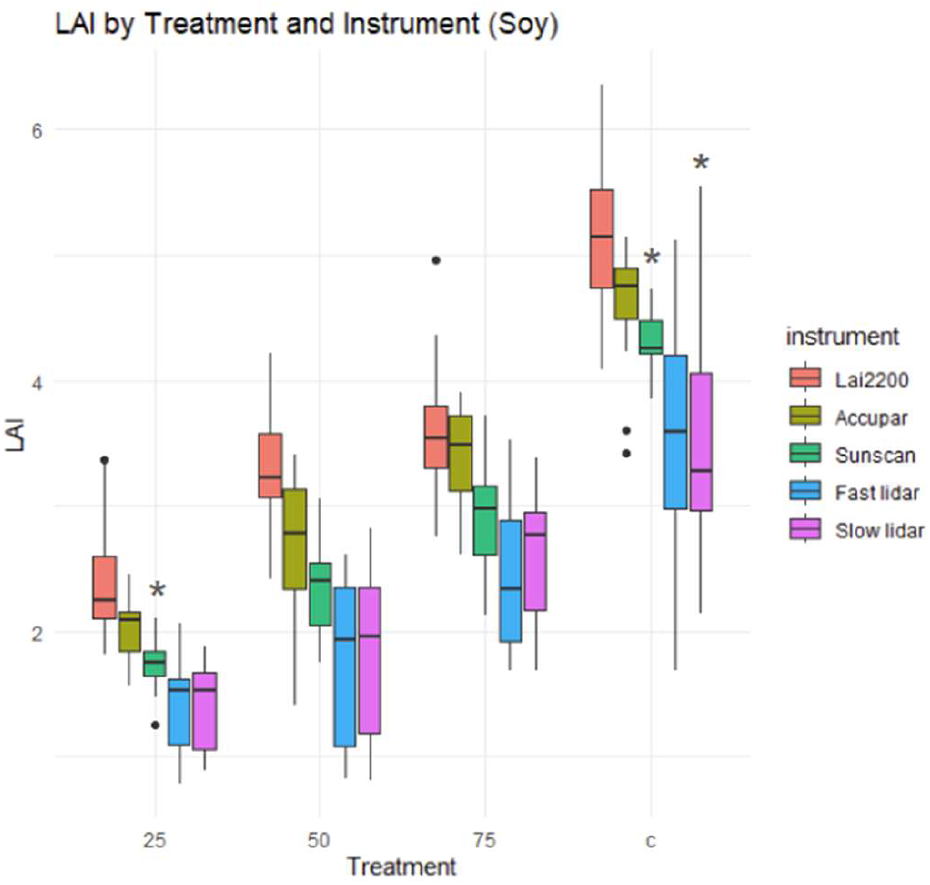
Distributions of LAI measurements in soybean for each method—AccuPAR, LAI-2200, SunScan, and fast and slow LiDAR—across thinning treatment levels: control, 75%, 50%, and 25% of initial plant density. Each boxplot includes all repeated measurements for a given instrument within a treatment. At each treatment level, variability for each method was compared to that of the LAI-2200 using a two-tailed F-test. Significant differences in variance are indicated with an asterisk (*p* < 0.05).

## Discussion

In corn, LiDAR-derived stand counts correlated well with manual counts, though slight underestimation was observed in denser plots. Fast-paced LiDAR scans were as precise as manual counts, while slow scans were even more precise at lower densities. Both methods had limitations. Manual counts may have lacked precision due to non-random placement of the meter stick, as people making the measurements may have avoided sparse areas, and only one meter of row is evaluated. By contrast, LiDAR scans cover the full plot length, which could increase precision. LiDAR accuracy, however, depended on scan quality. In both fast and slow scans, one outlier plot showed an undercount of two plants per square meter. Visual inspection of the point cloud revealed fallen leaves obscuring stalks, causing the algorithm to incorrectly merge several plants into one.

Bias between methods is not inherently problematic, especially when comparing treatments, since consistent bias cancels out when calculating treatment differences. Thus, without a true reference standard, one method is not necessarily “correct.” Nonetheless, it is informative to consider why two methods might differ and assess whether the causes are meaningful. Here, manual counts within the bounded meter stick were likely very accurate; variability may stem from where the measurement stick was placed but because plot density was largely uniform and the measurement protocol is relatively simple, any human bias was likely small. As such, the manual count may be considered the best available “true” value. LiDAR undercounting in dense plots suggests reduced detection accuracy under those conditions. Still, for many researchers, the reduced measurement time for LiDAR may outweigh these small differences, particularly when relative comparisons between plots are sufficient. Additionally, LiDAR scans can generate multiple traits (e.g., plant height, volume, LAI) from a single pass.

In soybean, LiDAR-based stand count estimations were unsuccessful using our system. The canopy was too dense for the persistent homology algorithm to resolve individual plant peaks. Although variability was low, this reflected consistent overestimation rather than true precision—the algorithm produced similar counts regardless of actual plant number. Scans were conducted during early pod development, when the canopy was particularly dense. However, early in the season (e.g., post-emergence), or late in the season when leaves have senesced, soybean canopies may be sparse enough for the algorithm to work effectively. Furthermore, higher resolution, multi-band LiDAR systems that include several measurement channels might help overcome this limitation.

Previous studies evaluated different LAI measurement approaches using ANOVA, comparing measurements to destructive sampling (Wilhelm et al., 2000). These analyses typically assess bias, assuming equal variances between instruments. However, precision is often more critical than bias. An instrument may systematically over-or underestimate LAI, and that may be acceptable provided the effect is consistent. In contrast, when other factors (e.g., cost, speed, ease of use) are equal, more precise tools increase statistical power and improve detection of treatment effects.

One obstacle to adopting new instruments is uncertainty about how their precision compares under specific conditions. Correlation analysis alone can be misleading, as variability may originate from either method. In this study, LAI correlation plots showed scatter, but variance analysis revealed that all non-LiDAR instruments had similar precision in corn, except at the highest LAI values where LiDAR precision decreased. Unlike correlation coefficients, variance estimates can be compared across studies. For example, if a new LiDAR-based LAI method is developed, its precision could be benchmarked against the values reported here. Though LiDAR was not the most precise method overall, it was by far the fastest, again offering a potential compromise between speed and precision.

Reduction in precision at high LAI and the inability to count plants in soybean canopies indicate a general problem using lidar in very dense canopies. This is likely caused by LiDAR beams being intercepted by the lower canopy, limiting penetration into the upper canopy. This results in either zero-gap errors during LAI estimation or the absence of distinct peaks for stand count. Improvements in LiDAR hardware could address these limitations. Multi-beam LiDARs scan at multiple angles simultaneously, enhancing canopy coverage. A wider field of view, capturing both front and rear of the device, would improve upper canopy penetration. Additionally, narrower beam widths would increase the likelihood of beams passing through small canopy gaps, enhancing both gap detection for LAI and point cloud resolution for stand count.

This study highlights the mixed performance of LiDAR in measuring stand count and LAI across corn and soybean. In soybean, LiDAR was ineffective due to dense canopies. In corn, it performed comparably to established methods. While precision decreased at high plant density, the dramatic increase in speed may justify its use in many applications. Moreover, LiDAR remains an evolving technology. With improved sensors and algorithms, it may become even more competitive. The three commercial canopy analyzers produced slightly different mean LAI values, as expected from design differences, but all showed consistent responses to changing plot density and comparable precision. Therefore, from a statistical power standpoint, these instruments are functionally equivalent. LiDAR, while fast and multifunctional, is currently limited by precision in dense canopies. However, technological advances in LiDAR may overcome these challenges and unlock broader use in high-throughput phenotyping.

## Acknowledgements

Funding for this work was provided by the DOE Center for Advanced Bioenergy and Bioproducts Innovation (U.S. Department of Energy, Office of Science, Office of Biological and Environmental Research under Award Number DE-SC0018420), by Gates Agricultural Innovations grant investment ID 57248 and the U.S. Department of Agriculture, Agricultural Research Service. Views expressed in this article are those of the authors and do not necessarily reflect those of the funding agencies acknowledged here

## Supplemental information

**Supplemental Figure 1.**
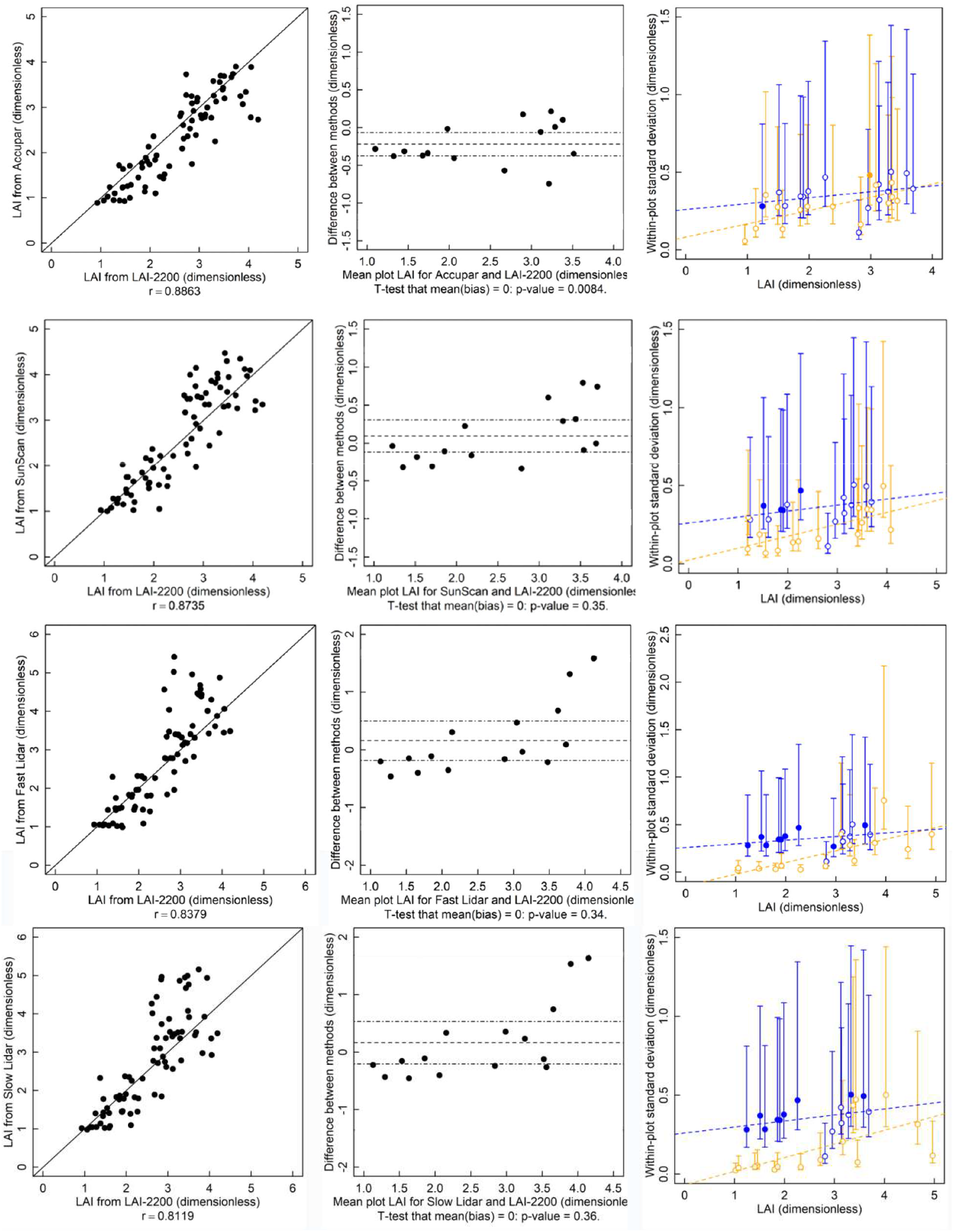
Correlation, bias and standard deviation of four methods measured in corn compared to the LAI-2200. Data were collected from 15 plots that were thinned 0, 25, 50 and 75 percent. Bias plots show the mean of five repeated measurements per method (dots), mean bias (dashed line), and 95% confidence interval of the bias (dashed-dotted line). The within-plot standard deviation of each method compares the LAI-2200 (blue symbols) with the alternative method (yellow symbol) with a two-tailed F-test. If a symbol is filled it means that the variance of that instrument is greater than other methods in the same plot (p < 0.05). Dashed lines are least square regressions that fit variance data but are for visualization only.

**Supplemental Figure 2.**
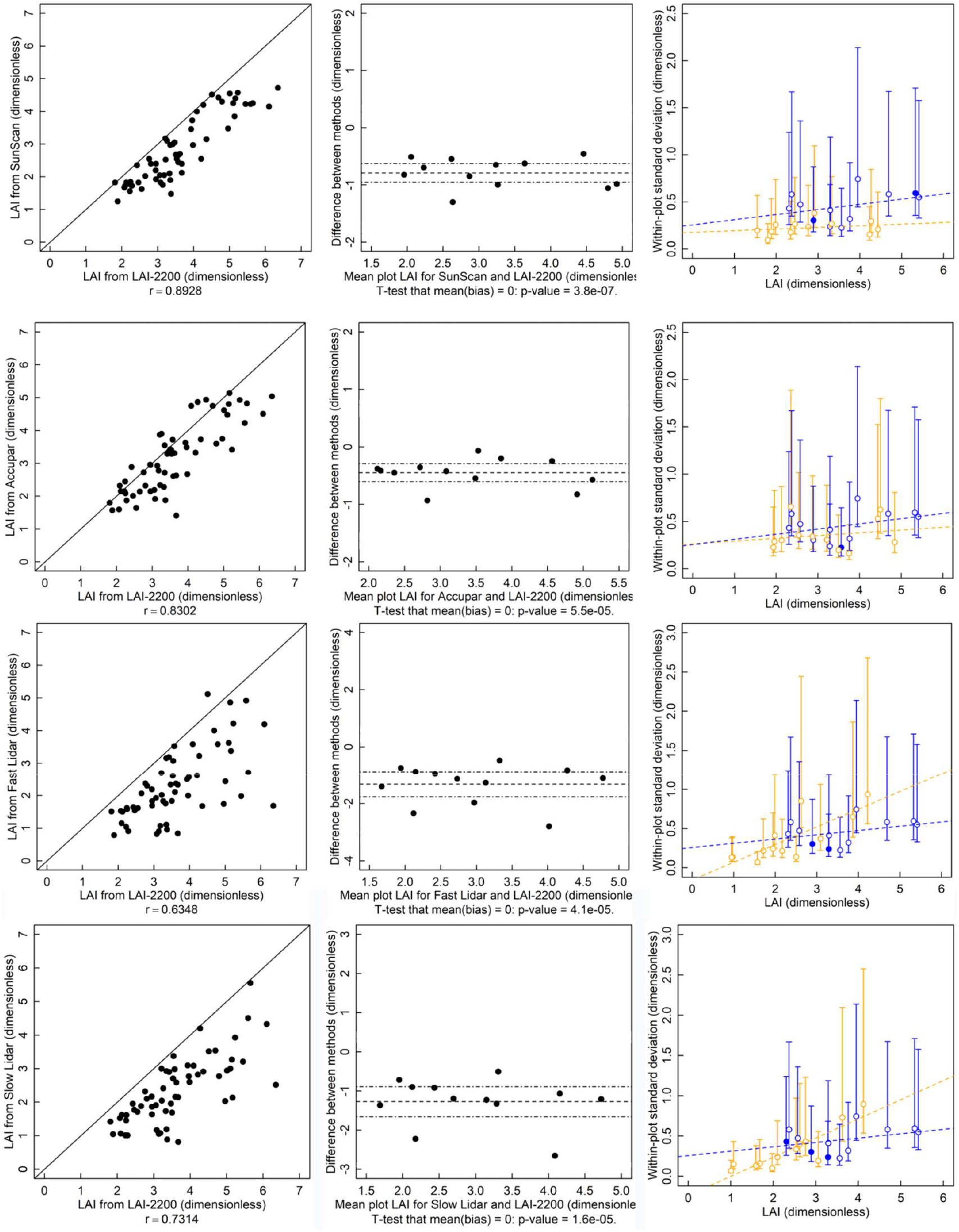
Correlation, bias and standard deviation of four methods measured in soybean compared to the LAI-2200. Data were collected from 15 corn plots that were thinned 0, 25, 50 and 75 percent. Bias plots show the mean of five repeated measurements per method (dots), mean bias (dashed line), and 95% confidence interval of the bias (dashed-dotted line). The within-plot standard deviation of each method compares the LAI-2200 (blue symbols) with the alternative method (yellow symbol) with a two-tailed F-test. If a symbol is filled it means that the variance of that instrument is greater than other methods in the same experimental plot (p < 0.05). Dashed lines are least square regressions that fit variance data but are for visualization only.

**Supplemental Figure 3.**
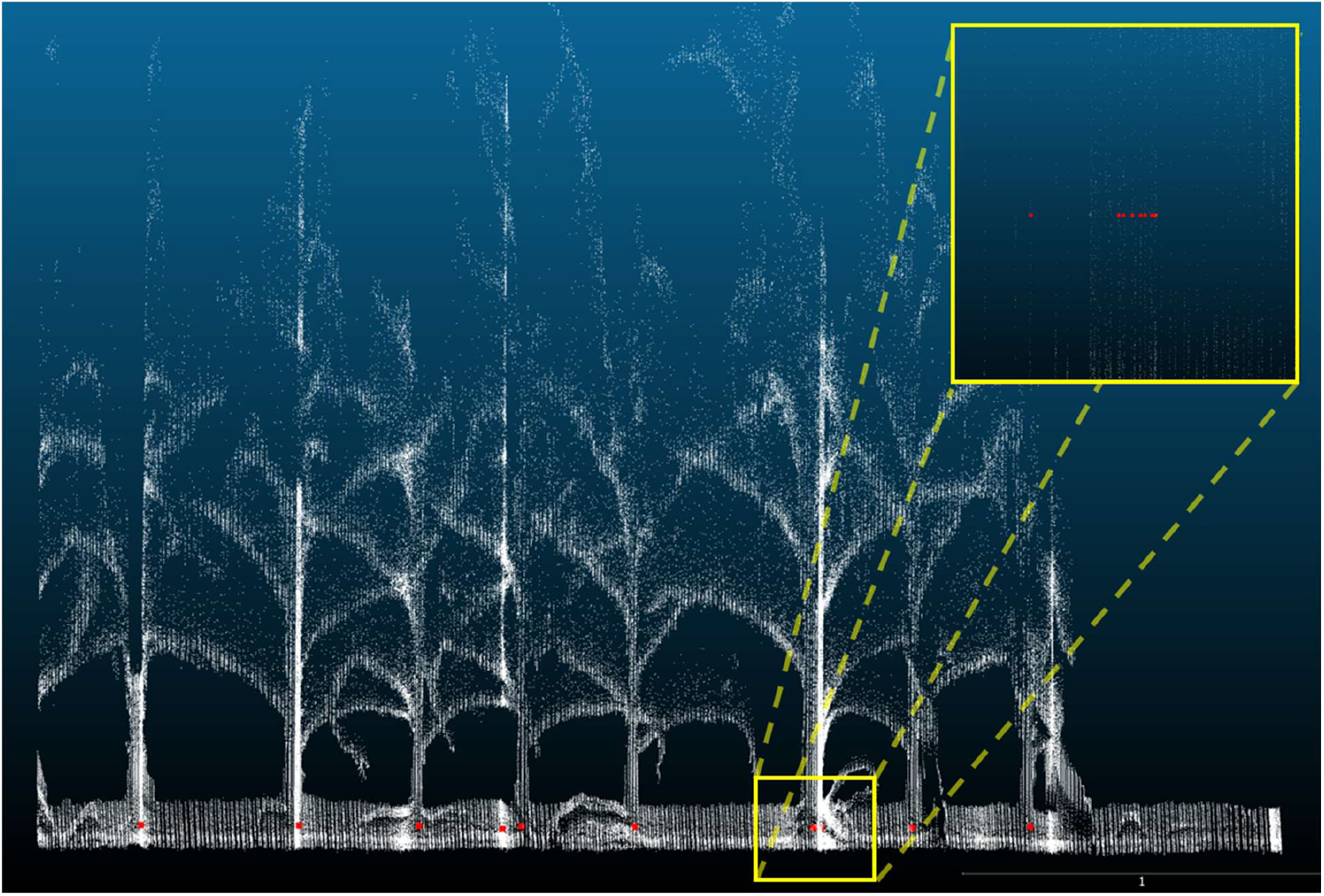
An example point cloud without z-filtering. The dense bands of scans compressed in the z-dimension were caused by the cart speed slowing. The lidar scans at a fixed frequency, and thus the lidar scanned the same area several times. The bands were counted multiple times by the persistence algorithm and were an issue obtaining accurate counts. The inset shows an enlarged view of the base of a plant that contains a dense patch of scans. The persistence algorithm counts that area eight times.

