## Supplemental Material for "Evaluation of LiDAR-based Canopy Trait Estimation in Midwestern Row Crops"

### Supplemental information

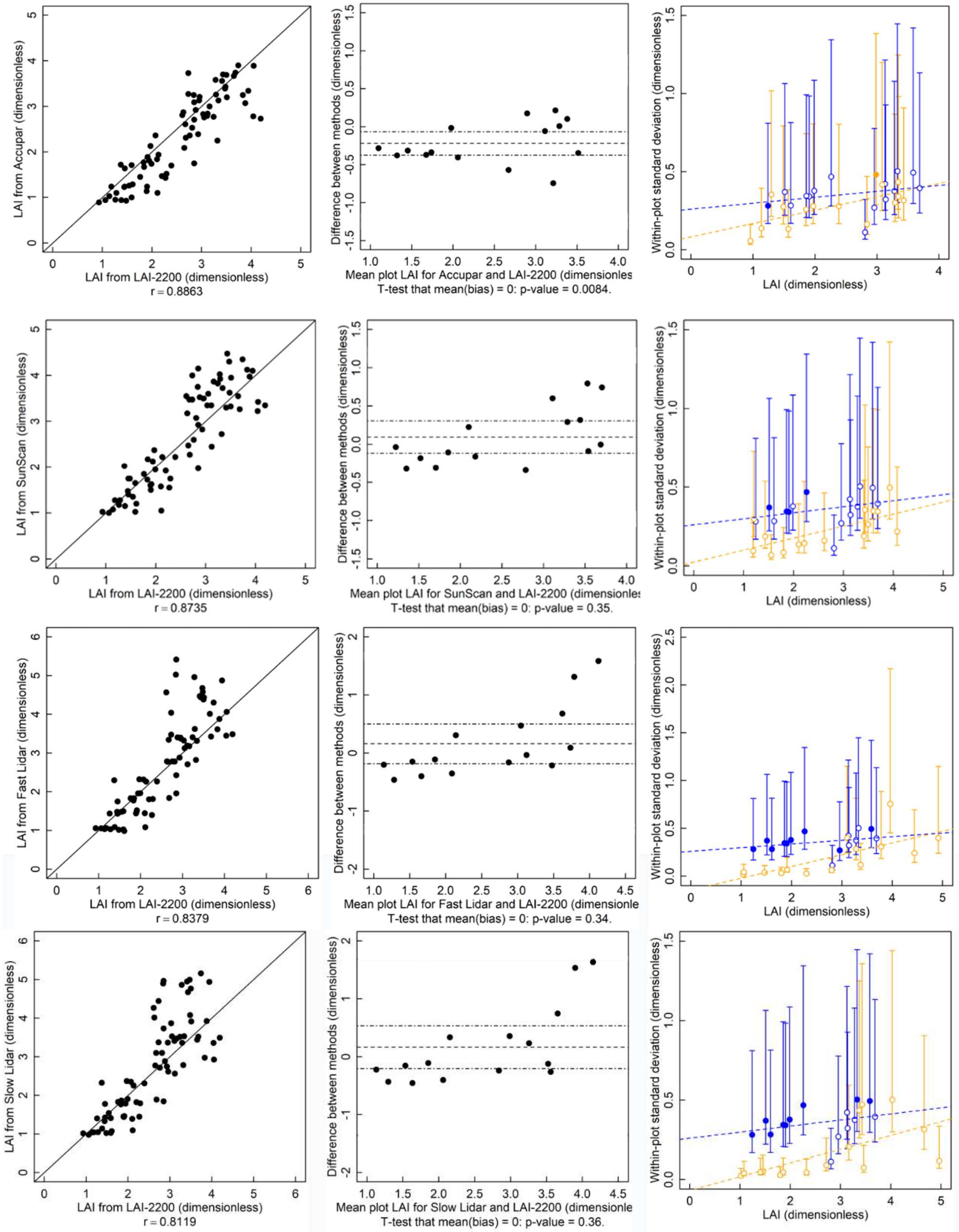


Supplemental Figure 1. Correlation, bias and standard deviation of four methods measured in corn compared to the LAI-2200. Data were collected from 15 plots that were thinned 0, 25, 50 and 75 percent. Bias plots show the mean of five repeated measurements per method (dots), mean bias (dashed line), and 95% confidence interval of the bias (dashed-dotted line). The within-plot standard deviation of each method compares the LAI-2200 (blue symbols) with the alternative method (yellow symbol) with a two-tailed F-test. If a symbol is filled it means that the variance of that instrument is greater than other methods in the same plot ($p < 0.05$). Dashed lines are least square regressions that fit variance data but are for visualization only.

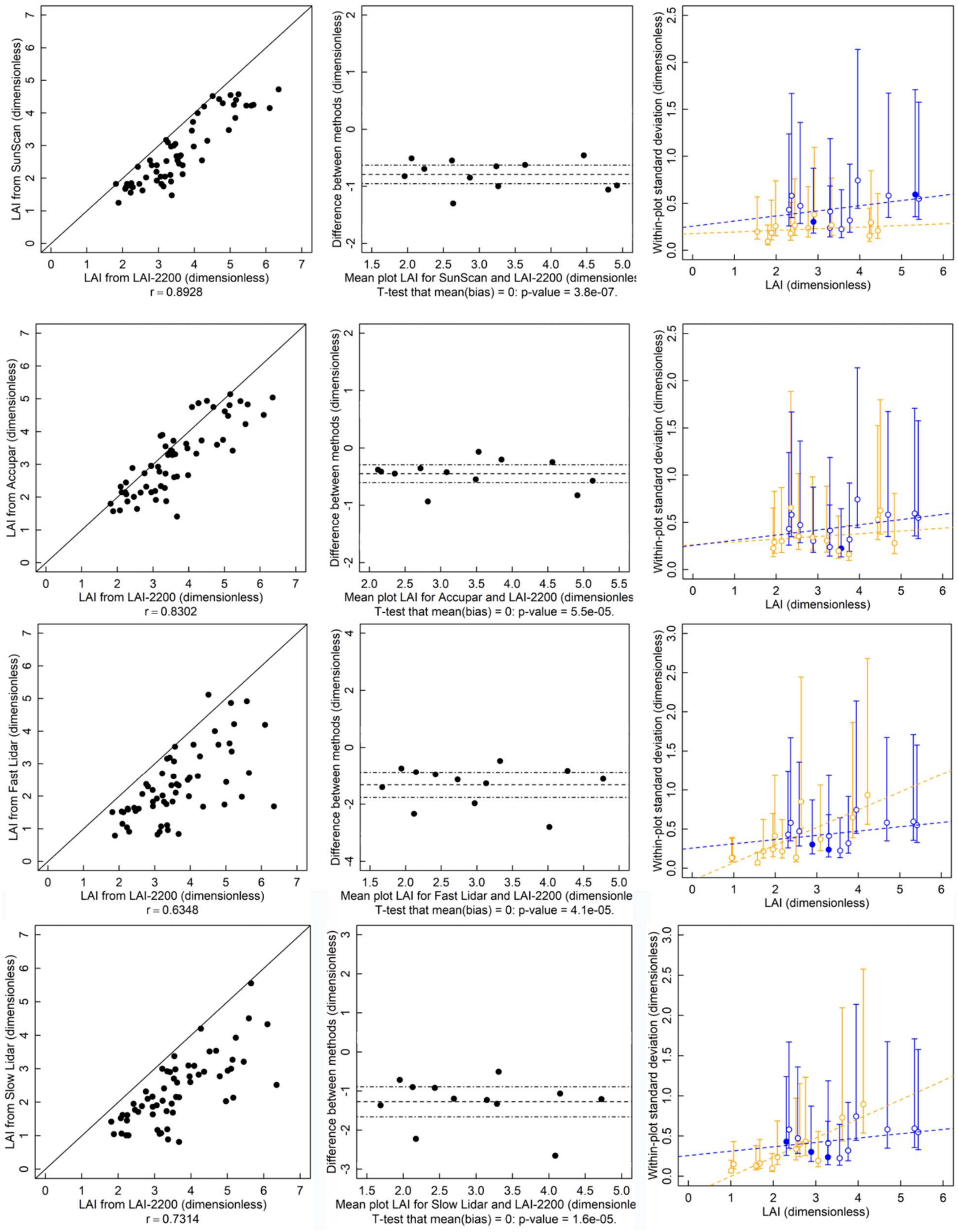


Supplemental Figure 2. Correlation, bias and standard deviation of four methods measured in soybean compared to the LAI-2200. Data were collected from 15 corn plots that were thinned 0, 25, 50 and 75 percent. Bias plots show the mean of five repeated measurements per method (dots), mean bias (dashed line), and 95% confidence interval of the bias (dashed-dotted line). The within-plot standard deviation of each method compares the LAI-2200 (blue symbols) with the alternative method (yellow symbol) with a two-tailed F-test. If a symbol is filled it means that the variance of that instrument is greater than other methods in the same experimental plot ($p < 0.05$). Dashed lines are least square regressions that fit variance data but are for visualization only.

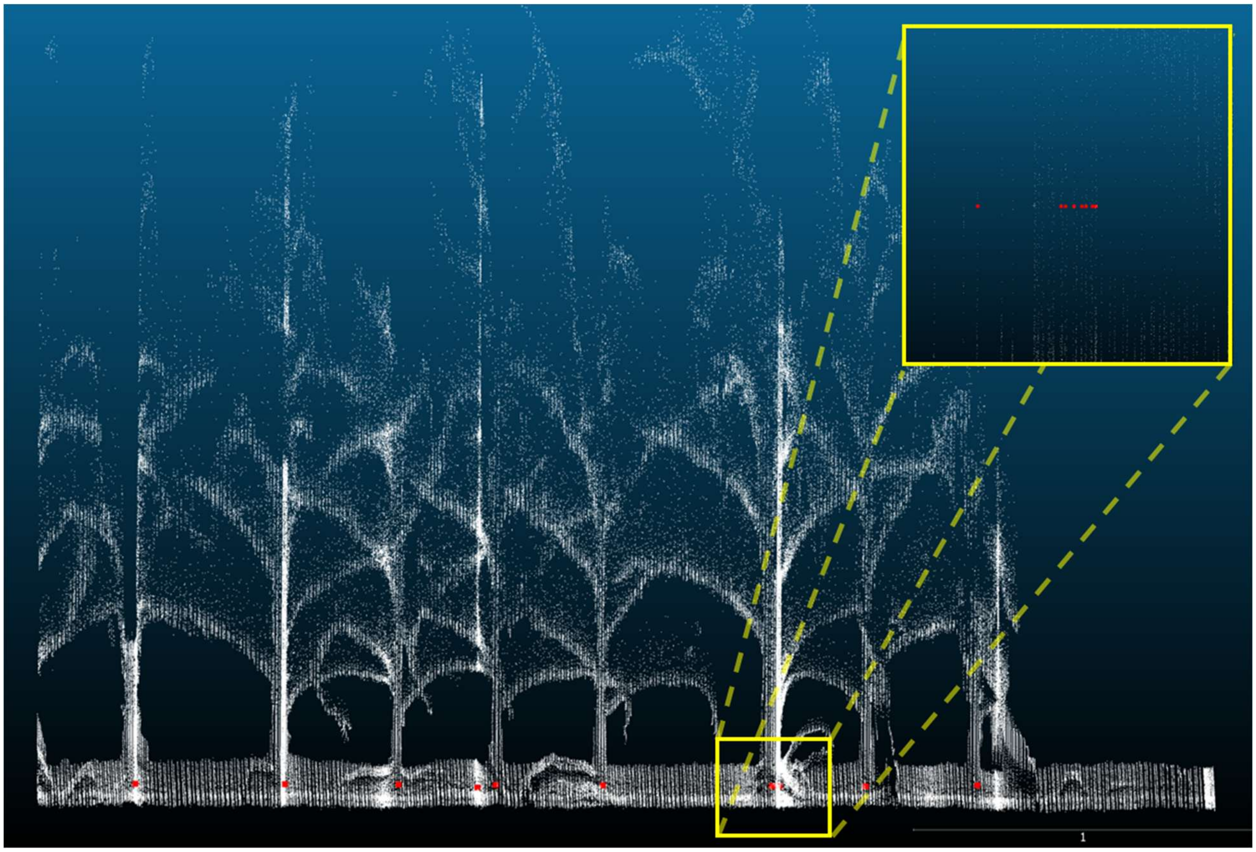


Supplemental Figure 3. An example point cloud without z-filtering. The dense bands of scans compressed in the z-dimension were caused by the cart speed slowing. The lidar scans at a fixed frequency, and thus the lidar scanned the same area several times. The bands were counted multiple times by the persistence algorithm and were an issue obtaining accurate counts. The inset shows an enlarged view of the base of a plant that contains a dense patch of scans. The persistence algorithm counts that area eight times.
